# Loss of choline agonism in the inner ear hair cell nicotinic acetylcholine receptor linked to the α10 subunit

**DOI:** 10.1101/2020.11.26.400291

**Authors:** Marcelo J. Moglie, Irina Marcovich, Jeremías Corradi, Agustín E. Carpaneto Freixas, Sofía Gallino, Paola V. Plazas, Cecilia Bouzat, Marcela Lipovsek, Ana Belén Elgoyhen

## Abstract

The α9α10 nicotinic acetylcholine receptor (nAChR) plays a fundamental role in inner ear physiology. It mediates synaptic transmission between efferent olivocochlear fibers that descend from the brainstem and hair cells of the auditory sensory epithelium. The α9 and α10 subunits have undergone a distinct evolutionary history within the family of nAChRs. Predominantly in mammalian vertebrates, the α9α10 receptor has accumulated changes at the protein level that may ultimately relate to the evolutionary history of the mammalian hearing organ. In the present work we investigated the responses of α9α10 nAChRs to choline, the metabolite of acetylcholine degradation at the synaptic cleft. Whereas choline is a full agonist of chicken α9α10 receptors it is a partial agonist of the rat receptor. Making use of the expression of α9α10 heterologous receptors, encompassing wild-type, heteromeric, homomeric, mutant, chimeric and hybrid receptors, and *in silico* molecular docking, we establish that the mammalian (rat) α10 nAChR subunit underscores the reduced efficacy of choline. Moreover, we show that whereas the complementary face of the α10 subunit does not play an important role in the activation of the receptor by ACh, it is strictly required for choline responses. Thus, we propose that the evolutionary changes acquired in the mammalian α9α10 nAChR resulted in the loss of choline acting as a full agonist at the efferent synapse, without affecting the triggering of ACh responses. This may have accompanied the fine-tuning of hair cell post-synaptic responses to the high frequency activity of efferent medial olivocochlear fibers that modulate the cochlear amplifier.

**Contribution to the Field Statement:** In the inner ear of mammals, several evolutionary changes have occurred resulting in an expansion of the hearing range to higher sound frequencies. Fine tuning of cochlear synapses is required for sound enconding. The synapse between efferent olivocochlear fibers, that descend from the hindbrain, and sensory hair cells modulates sound amplification at the periphery and it has been proposed as a major player in the expansion of the hearing range. The α9α10 nicotinic acetylcholine receptor, which mediates synaptic neurotransmission at the efferent fiber-hair cell synapses, has accumulated a high number of amino acid substitutions in the mammalian lineage. We now show that these evolutionary acquired changes have led to a mammalian receptor with a lower efficacy for choline, the metabolite produced at the synaptic cleft by acetylcholine degradation. Making use of molecular, electrophysiological and *in silico* simulations techniques we show that it is the α10 subunit the one responsible for the loss of full choline agonism on the efferent receptor in mammals. This functional change may prove fundamental to faithfully reproduce the high frequency activity of efferent medial olivocochlear fibers and the modulation of the cochlear amplifier.

## Introduction

The α9α10 nicotinic acetylcholine receptor (nAChR) belongs to the pentameric family of ligand-gated ion channels (Elgoyhen et al., 1994; Elgoyhen et al., 2001; Sgard et al., 2002). Each subunit has a large extracellular N-terminal region, four transmembrane helices (M1–M4), and an intracellular domain. At the interface of the extracellular domains of adjacent subunits lies the orthosteric binding site, composed of a principal component or face provided by one subunit, which contributes three loops of highly conserved residues (loops A–C), and a complementary component of the adjacent subunit, which contributes three loops (loops D–F) with lower levels of sequence conservation among subunits (Thompson et al., 2010). Conserved aromatic residues present within the loops participate in cation-π interactions with the agonists that are fundamental for triggering receptor gating (Karlin, 2002).

The α9α10 receptor is an atypical member of the nAChR family. Both the α9 and α10 subunits have low amino acid sequence identity when compared to other member subunits (Elgoyhen et al., 1994; Elgoyhen et al., 2001; Franchini and Elgoyhen, 2006; Elgoyhen et al., 2009; Lipovsek et al., 2012; Lipovsek et al., 2014; Marcovich et al., 2020) and α9α10 receptors have a baroque pharmacological profile (Rothlin et al., 1999; Verbitsky et al., 2000; Rothlin et al., 2003; Ballestero et al., 2005). Indeed, nicotine, the canonical agonist that characterizes the receptor subfamily, does not activate α9α10 nAChRs, but blocks ACh-gated currents (Elgoyhen et al., 1994; Elgoyhen et al., 2001; Sgard et al., 2002). Moreover, the α9α10 nAChR has a mixed nicotinic and muscarinic profile, since it is blocked by the nicotinic antagonists curare and α-bungarotoxin and the muscarinic antagonist atropine (Elgoyhen et al., 1994; Elgoyhen et al., 2001; Sgard et al., 2002). In addition, the α9α10 receptor shares pharmacological properties with type A γ-aminobutyric acid, glycine, and type 3 serotonin receptors, also members of the pentameric family of ligand-gated ion channels (Rothlin et al., 1999; Rothlin et al., 2003).

The atypical features of the α9α10 receptor prompted the hypothesis that α9 and α10 subunits have undergone a distinct evolutionary history within the family of nAChRs. Using codon-based likelihood models we showed that mammalian, unlike non-mammalian, α10 subunits have been under selective pressure and acquired a greater than expected number of non-synonymous amino acid substitutions in their coding region (Franchini and Elgoyhen, 2006; Elgoyhen and Franchini, 2011). Moreover, mammalian specific amino acid substitutions in the α9 subunit, that show an increased posterior probability of functional divergence in this clade, are involved in the higher relative calcium permeability of mammalian α9α10 receptors (Lipovsek et al., 2014; Marcovich et al., 2020). Overall, these patterns of evolutionary changes at the protein level may relate to the evolutionary history of the mammalian hearing organ (Marcovich et al., 2020), which has the highest frequency sensitivity among vertebrate auditory systems (Manley, 2000). In particular, α9α10 nAChRs mediate the synapses between efferent fibers and sensory hair cells that modulate sound amplification processes, including the mammalian-exclusive prestin-driven somatic electromotility of outer hair cells (OHCs) (Katz and Elgoyhen, 2014; Goutman et al., 2015). Overall, the distinct evolutionary history of mammalian α9α10 nAChRs resulted in differential calcium permeability, current-voltage relationship and desensitization profile of α9α10 receptors across vertebrate species (Lipovsek et al., 2012; Lipovsek et al., 2014; Marcovich et al., 2020), together with the loss of functional homomeric α10 receptors (Elgoyhen et al., 2001; Sgard et al., 2002; Lipovsek et al., 2012) and a non-equivalent contribution of different subunit interfaces to functional binding sites in mammals (Boffi et al., 2017).

One striking feature of the pharmacology of mammalian α9α10 receptors, compared to other nAChRs, is the scarcity of identified compounds capable of behaving as agonists of the receptor, since most typical nicotinic agonists block rat α9α10 nAChRs (Verbitsky et al., 2000; Elgoyhen et al., 2001). Whether these observations are related to the peculiar evolutionary history of the mammalian receptor, and do not therefore extend to other non-mammalian α9α10 nAChRs, is still an open question. In the present work we have addressed this by comparing the effects of classical nicotinic agonists: nicotine, carbachol, DMPP and choline, on rat and chicken α9α10 nAChRs. We report that, as for rat receptors, nicotine does not activate but blocks chicken α9α10 nAChRs. However, whereas choline is a partial agonist of rat α9α10 receptors, it is a full agonist of chicken α9α10 receptors. Similarly, DMPP has a higher efficacy in chicken receptors. Using hybrid and chimeric receptors we show that the lower agonistic efficacy of choline in rat α9α10 receptors is linked to the extracellular region of the mammalian α10 subunit. Most importantly, we describe that complementary components of the ligand-binding site provided by the rat α10 subunit non-equivalently contribute to receptor activation by ACh and choline. Thus, whereas ACh does not utilize interfaces where α10 provides the complementary component to elicit maximal responses, choline requires fully competent α10 interfaces for receptor activation, suggesting a requirement for a higher degree of ligand occupancy when choline is the agonist. In line with these results, molecular docking simulations indicate that choline binds at all interfaces with a different orientation with respect to that of ACh, and it does so with differing frequencies depending on which subunit contributes the complementary side. Overall, we propose that the loss of choline full agonism in mammalian α9α10 receptors was driven by changes in the α10 subunit. This may have resulted from functional selection pressure on the fine-tuning of cholinergic responses within the mammalian efferent synapse.

## Materials and Methods

### Expression of Recombinant Receptors in *X. laevis* oocytes

For expression studies rat and chicken α9 and α10 nAChR subunits subcloned into a modified pGEMHE vector were used. Capped cRNAs were *in vitro* transcribed from linearized plasmid DNA templates using RiboMAX™ Large Scale RNA Production System (Promega, Madison, WI, USA). The maintenance of *X. laevis* and the preparation and cRNA injection of stage V and VI oocytes have been described in detail elsewhere (Verbitsky et al., 2000). Typically, oocytes were injected with 50 nl of RNase-free water containing 0.01 to 1.0 ng of cRNA (at a 1:1 molar ratio for heteromeric receptors) and maintained in Barth’s solution (in mM: NaCl 88, Ca(NO_3_)_2_ 0.33, CaCl_2_ 0.41, KCl 1, MgSO_4_ 0.82, NaHCO_3_ 2.4, HEPES 10) at 18°C. Electrophysiological recordings were performed 2 to 6 days after cRNA injection under two-electrode voltage clamp with an Oocyte Clamp OC-725B or C amplifier (Warner Instruments Corp., Hamden, CT, USA). Recordings were filtered at a corner frequency of 10 Hz using a 900BT Tunable Active Filter (Frequency Devices Inc., Ottawa, IL, USA). Data acquisition was performed using a Patch Panel PP-50 LAB/1 interphase (Warner Instruments Corp., Hamden, CT, USA) at a rate of 10 points per second. Both voltage and current electrodes were filled with 3 M KCl and had resistances of ~1 MΩ. Data were analyzed using Clampfit from the pClamp 6.1 software. During electrophysiological recordings, oocytes were continuously superfused (~15 ml/min) with normal frog saline composed of: 115 mM NaCl, 2.5 mM KCl, 1.8 mM CaCl_2_, and 10 mM HEPES buffer, pH 7.2. Drugs were added to the perfusion solution for application. The membrane potential was clamped to −70 mV. To minimize activation of the endogenous Ca^2+^ sensitive chloride current (Elgoyhen et al., 2001), all experiments were performed in oocytes incubated with the Ca^2+^ chelator 1,2-bis (2-aminophenoxy)ethane-N,N,N’,N’-tetraacetic acid-acetoxymethyl ester (BAPTA-AM, 100 μM) for 3h before electrophysiological recordings. Concentration-response curves were normalized to the maximal ACh response in each oocyte. For the inhibition curves, nicotine was added to the perfusion solution for 2 min before the addition of 10 μM ACh and then coapplied with this agonist and responses were referred to as a percentage of the response to ACh. The mean and S.E.M. of peak current responses are presented. Agonist concentration-response curves were iteratively fitted, using Prism 6 software (GraphPad Software Inc., La Jolla, CA, USA), with the equation: I/Imax = A^nH^/(A^nH^ + EC_50_^nH^), where I is the peak inward current evoked by agonist at concentration A; Imax is the current evoked by the concentration of agonist eliciting a maximal response; EC_50_ is the concentration of agonist inducing half-maximal current response, and nH is the Hill coefficient. An equation of the same form was used to analyze the concentration dependence of antagonist induced blockage. The parameters derived were the concentration of antagonist producing a 50% block of the control response to ACh (IC_50_) and the associated interaction coefficient (*n*H).

The α10W55T mutant subunit was produced as previously described (Boffi et al., 2017) using Quick change XL II kit (Stratagene, La Jolla, CA, USA). The chimeric α10x construct was generated in two steps. First, the DNA fragment encoding the α10 N-terminal domain was replaced by the DNA sequence encoding the α9 N-terminal domain, by overlap-extension PCR (Horton et al., 1989). Briefly, the DNA sequence of the α9 N-terminal domain was amplified from an α9 pGEMHE construct using primers 5’GGGCGAATTAATTCGAGCTC3’ and 5’CACCTTCACTCTCCTTCTGAAGCGCCGCGCTGCAGCCTACGTGTG3’, and the DNA sequence of the α10 subunit from TM1 to the C-terminal domain was amplified from an α10 pSGEM construct using primers 5’CACACGTAGGCTGCAGCGCGGCGCTTCAGAAGGAGAGTGAAGGTG3’ and 5’GCTATGACCATGATTACGCC3’. The chimeric subunit was amplified using the DNA fragments generated in the two PCRs described above and primers 5’GGGCGAATTAATTCGAGCTC3’ and 5’GCTATGACCATGATTACGCC3’, and subcloned in pSGEM vector. In a second step, conversion of both the pre TM1 and the TM2-TM3 loop from the chimera to the corresponding α9 sequence was performed by QuikChange Multi Site-Directed Mutagenesis kit (Stratagene) using primers 5’GGCATGCTCTCGGCCACCATCAGCTGGAAGACGG3’ and 5’GGGCAGCAGGAGGTTGACGATGTAGAATGAAGAGCGGCGCTTCAG3’. All mutant and chimeric subunits were confirmed by sequencing.

### Recordings from hair cells

Chicken auditory organ (basilar papilla) was dissected from the temporal bone of embryonic chickens (white Leghorns either sex, 17–20 d *in ovo*). The tegmentum vasculosum was removed and the tectorial membrane was stripped from the basilar papilla with most of the short hair cells (SHCs) still attached. This “tectorial preparation” was then inverted and secured in the recording chamber by spring clips. SHCs were recorded in a region 25–50% the distance from the apical (lagenar) to the basal tip of the basilar papilla of the chicken. Apical turns of the organ of Corti were excised from Balb/C mice of either sex between postnatal day 11 (P11) and P13, (around the onset of hearing in altricial rodents). At this age outer hair cells (OHCs) are already innervated by the MOC fibers (Pujol et al., 1998; Simmons, 2002). The tectorial membrane was removed and the organ of Corti was positioned under an insect pin affixed to a cover-slip with a drop of Sylgard.

Basilar papilla and cochlear preparations were placed into a chamber on the stage of an upright microscope (Olympus BX51WI, Center Valley, PA, USA) at room temperature and used within 2 h. Cells were visualized on a monitor via a water immersion objective (60× magnification), differential interference contrast optics, and a CCD camera (Andor iXon 885, Belfast, UK). The preparation was superfused continuously at 2–3 mL per min with extracellular saline solution of an ionic composition similar to that of the perilymph: 144 mM NaCl, 5.8 mM KCl, 1.3 mM CaCl_2_, 0.9 mM MgCl_2_, 0.7 mM NaH_2_PO_4_, 5.6 mM D-glucose and 10 mM HEPES buffer (pH 7.4). Solutions containing ACh or choline were prepared in this same saline and delivered by a gravity-fed multichannel glass pipette (150-μm tip diameter).

Whole-cell, tight-seal voltage-clamp recordings were made with 1 mm borosilicate glass micropipettes (WPI, Sarasota, FL, USA) ranging from 6 to 8 MΩ tip resistance. Series resistance errors were not compensated for. The recording pipette contained the following (in mM): 140 KCl, 3.5 MgCl_2_, 2 CaCl_2_, 5 EGTA, 5 HEPES, 5 mM phosphocreatine-Na_2_ and 2.5 Na_2_ATP, titrated to pH 7.2 with KOH. Osmolarity was adjusted to 295 mOsM. All recordings were performed at room temperature (22–25 °C).

Electrophysiological recordings were performed using a Multiclamp 700B amplifier (Molecular Devices, San Jose, CA, USA), low-pass-filtered at 2–10 kHz, and digitized at 50 kHz via a National Instruments board. Data were acquired using WinWCP (J. Dempster, University of Strathclyde, Glasglow, Scotland). Recordings were analyzed with custom-written routines in IgorPro 6.37 (Wavemetrics, Lake Oswego, OR, USA).

### Molecular modeling and docking

Homology models of the extracellular domain of the chick and rat α9α10 nAChRs were as in (Boffi et al., 2017). They were created with SWISS MODEL (Schwede et al., 2003; Arnold et al., 2006; Bordoli et al., 2009) using the monomeric structure of the human α9 subunit as the template (Protein Data Bank ID 4UY2;(Zouridakis et al., 2014)). The monomeric models of these proteins were then structurally aligned to the pentameric structure of *Lymnaea stagnalis* ACh binding protein (AChBP) bound to ACh (Protein Data Bank ID 3WIP; (Olsen et al., 2014)) using the program STAMP (Russell and Barton, 1992) from visual molecular dynamics (Humphrey et al., 1996) to obtain pentameric models with a (α9)_2_(α10)_3_ stoichiometry bound to ACh. Four different types of possible binding site interfaces were included: α9/α9, α9/α10, α10/α9 and α10/α10. For each interface, the first subunit provides the principal face and the second provides the complementary face. The models were energy minimized to relax steric clashes using spdbviewer (Guex and Peitsch, 1997), and were used for docking studies after deletion of ACh from the models. Using Autodock 4.2 (Morris et al., 2009) choline and ACh were downloaded from PubChem and docked into each of the four types of interfaces for rat and chicken subunits. One hundred genetic algorithm runs were performed for each condition. Residues R57, R111, and R117 were set as flexible to avoid steric and/or electrostatic effects that may impair agonist docking into the binding site (Boffi et al., 2017). Clustering of the results was done with Autodock based on a root-mean-square deviation cutoff of 2.0 Å. Docking results were corroborated in three different procedures. The most representative docking result was plotted with Discovery Studio Visualizer 4.5 (Dessault Systèmes BIOVIA, San Diego, 2016).

### Statistical Analysis

Statistical analysis was performed using the software Infostat (Universidad Nacional de Córdoba). The non-parametric Mann-Whitney test was used to perform comparisons between two groups and Kruskal-Wallis one-way analysis of variance followed by Conover’s test for comparisons between multiple groups. Friedman’s test was used for comparisons between paired samples. Differences between samples were considered significant when p < 0.05. All raw data and analysis code are available upon request.

### Materials

All drugs were obtained from Sigma-Aldrich (St. Louis, MO, USA). ACh, choline, DMPP, carbachol and nicotine were dissolved in distilled water as stocks and stored aliquoted at −20°C. BAPTA-AM was stored at −20°C as aliquots of a 100 mM solution in dimethyl sulfoxide (DMSO), thawed and diluted 1000-fold into Barth’s solution shortly before incubation of the oocytes.

All experimental protocols were carried out in accordance with the National Institute of Health Guide for the Care and Use of Laboratory Animals (NIH Publications No. 80-23) revised 1978 and INGEBI Institutional Animal Care and Use Committee.

## Results

### Cholinergic agonists show lower efficacy on rat α9α10 receptors

The unique pharmacological profile of mammalian α9α10 receptors (Rothlin et al., 1999; Verbitsky et al., 2000; Rothlin et al., 2003), and the observation that mammalian α10 subunits have been under selection pressure (Franchini and Elgoyhen, 2006; Elgoyhen and Franchini, 2011), prompted us to examine whether this baroque pharmacology is recapitulated in receptors from a non-mammalian species. Nicotine, the prototypic agonist of nAChRs, is an antagonist of mammalian α9α10 receptors (Elgoyhen et al., 1994; Elgoyhen et al., 2001; Sgard et al., 2002), and did not elicit currents in oocytes expressing either rat or chicken α9α10 receptors. Instead, nicotine acted as an antagonist of both receptors (Fig. 1A, Table 1 and (Elgoyhen et al., 2001)), blocking responses to 10 μM ACh in a concentration-dependent manner (rat IC_50_ = 46.3 ± 12.4 μM; n=6; chicken IC_50_ = 39.4 ± 3.4 μM; n=6; p=0.81, Mann-Whitney test).

**Table 1.**
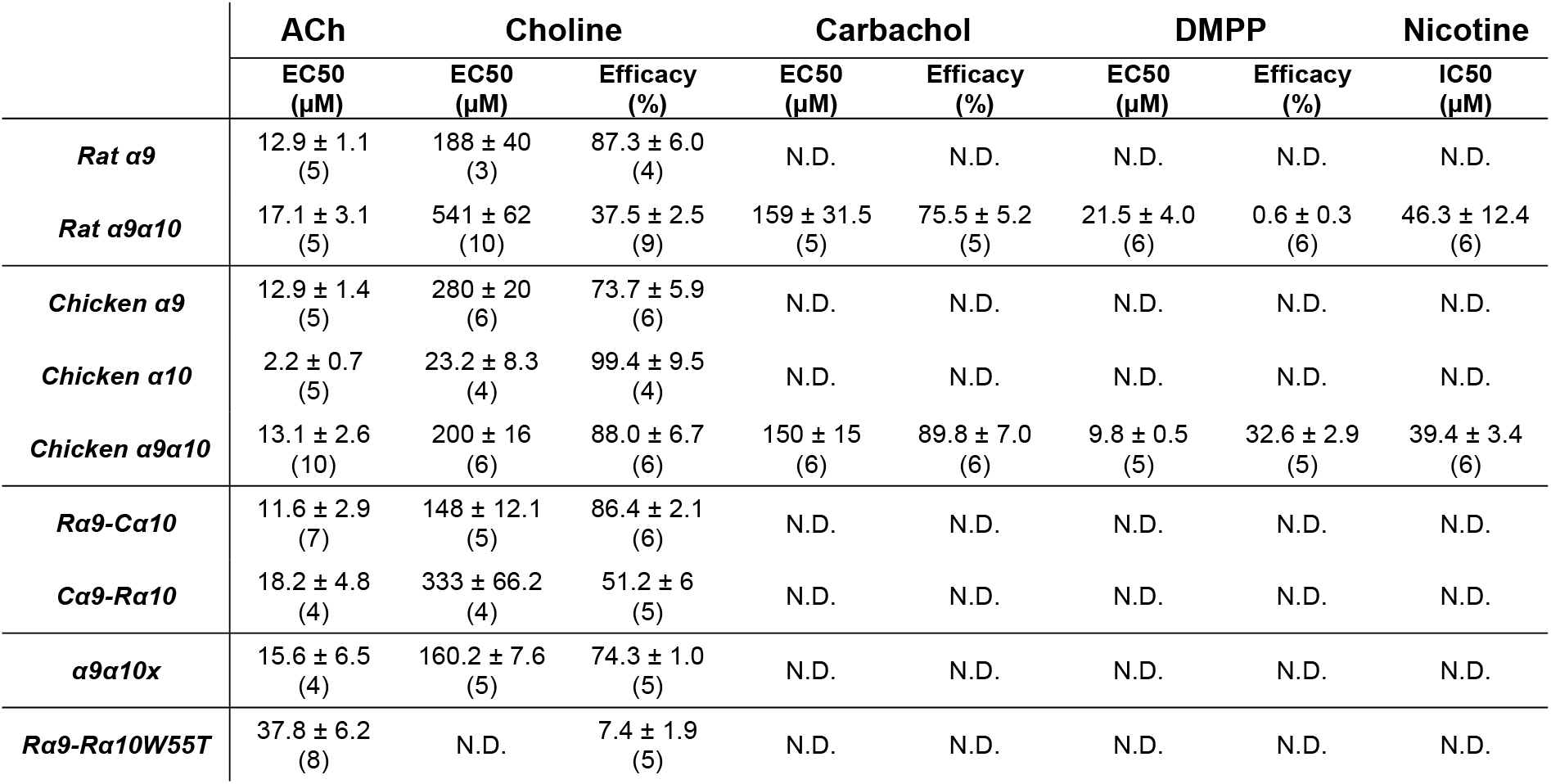
Estimated values of EC_50_ and efficacy for agonists of α9α10 receptors and IC_50_ value for the antagonist nicotine. Values are mean ± S.E.M.. Between brackets, number of oocytes per experiment. R, rat. C, chicken. N.D., not determined.

**Figure 1.**
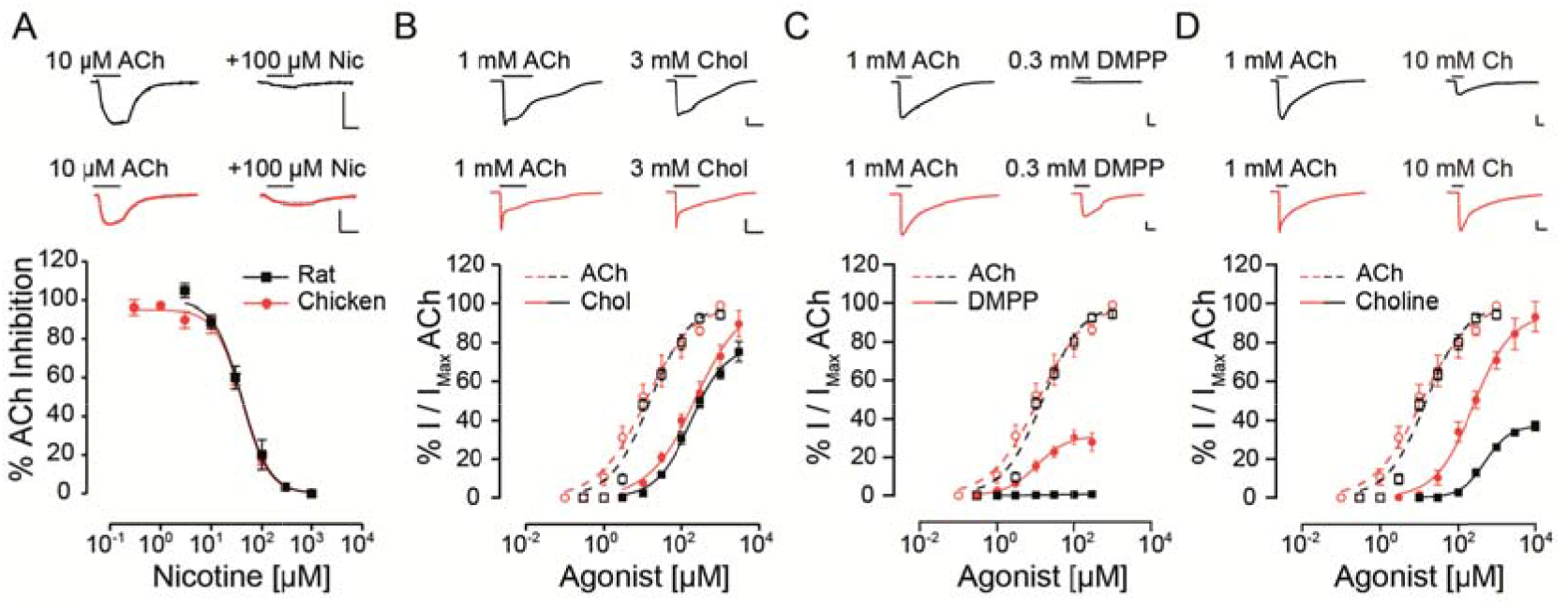
Responses of rat and chicken recombinant heteromeric α9α10 nAChRs to nicotinic agonists. A. Top panel: representative traces of responses to 10 μM ACh (left) blocked by co-application of 100 μM nicotine (right) for rat (black traces) and chicken (red traces) α9α10 receptors. Bottom panel: inhibition curves for nicotine for rat and chicken α9α10 receptors. Responses to 10 μM ACh, co-applied with increasing concentrations of nicotine were normalized to the responses to 10 μM ACh alone. Values are mean ± S.E.M. for 6-7 oocytes. B-D. Top panels: representative maximal responses to ACh and carbachol (B), DMPP (C) and chol ne (D) for rat and chicken α9α10 receptors. Bottom panels: concentration-response curves for ACh (dotted lines) and carbachol (B), DMPP (C) and choline (D) (solid lines) for rat (black) and chicken (red) α9α10 receptors. Values are normalized to the maximal response to ACh obtained in each oocyte. Values are mean ± S.E.M. for 5-9 oocytes.

Next, we analyzed the effect of different nAChR agonists on α9α10 receptors from the two studied species. Previous work showed that ACh elicits maximal responses in both rat and chicken α9α10 receptors, with similar near 10 μM EC_50_ values (Elgoyhen et al., 2001; Lipovsek et al., 2012). The cholinergic agonist carbachol elicited concentration-dependent responses in oocytes expressing α9α10 receptors (Fig. 1B and Table 1) with maximal responses that were 75 ± 5 % (rat, n=5) and 89 ± 7 % (chicken, n=6) of the maximum response elicited by 1 mM ACh (p=0.93, Mann-Whitney test). EC_50_ values for carbachol were similar for the receptors from both species: 159 ± 31.5 μM (rat; n=5), 150 ± 15 μM (chicken; n=6); p=0.42, Mann-Whitney test. The potency of carbachol was therefore nearly 12 times lower compared to that of ACh both in rat and chicken α9α10 recombinant receptors (Elgoyhen et al., 2001; Lipovsek et al., 2012). As previously reported (Elgoyhen et al., 2001), the nicotinic agonist DMPP behaved as a very weak partial agonist of rat α9α10 nAChRs (Fig. 1C and Table 1), with a maximal response that was 0.6 ± 0.3 %, (n=6) of that observed with 1 mM ACh (n=6). Surprisingly, the efficacy of DMPP on chicken α9α10 receptors was significantly higher (p=0.004 Mann-Whitney test compared to rat), reaching 32.6 ± 2.9 % of that of ACh with an EC_50_ of 9.8 ± 0.5 μM (n=5; p=0.03 Mann-Whitney test, chick vs rat). Finally, choline behaved as a partial agonist on rat heteromeric α9α10 receptors (Fig. 1D and Table 1), with a maximal response that was 37 ± 3 % of that produced by 1 mM ACh and an EC_50_ of 541 ± 62 μM (n=10). As observed for DMPP, the efficacy of choline on chicken α9α10 receptors was higher than that observed for rat receptors (n=6; p=0.0004 Kruskal-Wallis test), reaching a maximum response of 88 ± 7 % of the maximum response to 1 mM ACh, thus behaving nearly as a full agonist. Additionally, the EC_50_ of 200 ± 16 μM (n=6) for choline in chicken α9α10 receptors was nearly three times lower than that observed for rat receptors (EC_50_= 541 ± 62 (n=10), p=0.0025 Kruskal-Wallis test). In summary, while inhibition by nicotine and agonism by carbachol were similar between chicken and rat receptors, choline and DMPP presented higher efficacies for the avian than for the mammalian α9α10 heteromeric nAChRs.

### Choline elicits near-maximal responses in chicken, but not in mouse, hair cells

Choline is naturally present at cholinergic synapses as a degradation product of ACh (Wehrwein et al., 2016) and its effect on nAChRs can have important implications for synaptic function (Albuquerque et al., 1998). The synapse between efferent fibers and hair cells is mediated by α9α10 nicotinic receptors coupled to the activation of SK2 calcium dependent potassium channels, resulting in an inhibitory response (Fuchs and Murrow, 1992; Dulon and Lenoir, 1996; Katz et al., 2004; Gomez-Casati et al., 2005; Elgoyhen and Katz, 2012). We analyzed the effect of choline in mouse and chicken hair cells in acute explants, in comparison to ACh responses. Figure 2A shows representative responses to both ACh and choline at a holding potential of −40 mV. These responses were outward, resulting from the activation of SK2 potassium channels coupled to the cholinergic responses. For both chicken and mouse hair cells we observed maximal responses to 3 mM ACh. However, and in agreement with that observed for recombinant α9α10 receptors, choline exhibited higher efficacy in chicken hair cells (90 ± 6 %, n=3) when compared to that on mouse hair cells (50 ± 9 %, n=6; p=0.024 Mann-Whitney test). These observations suggest that the differences in agonistic behavior identified in recombinant α9α10 receptors are recapitulated by the native receptors present in chicken and mouse hair cells.

**Figure 2.**
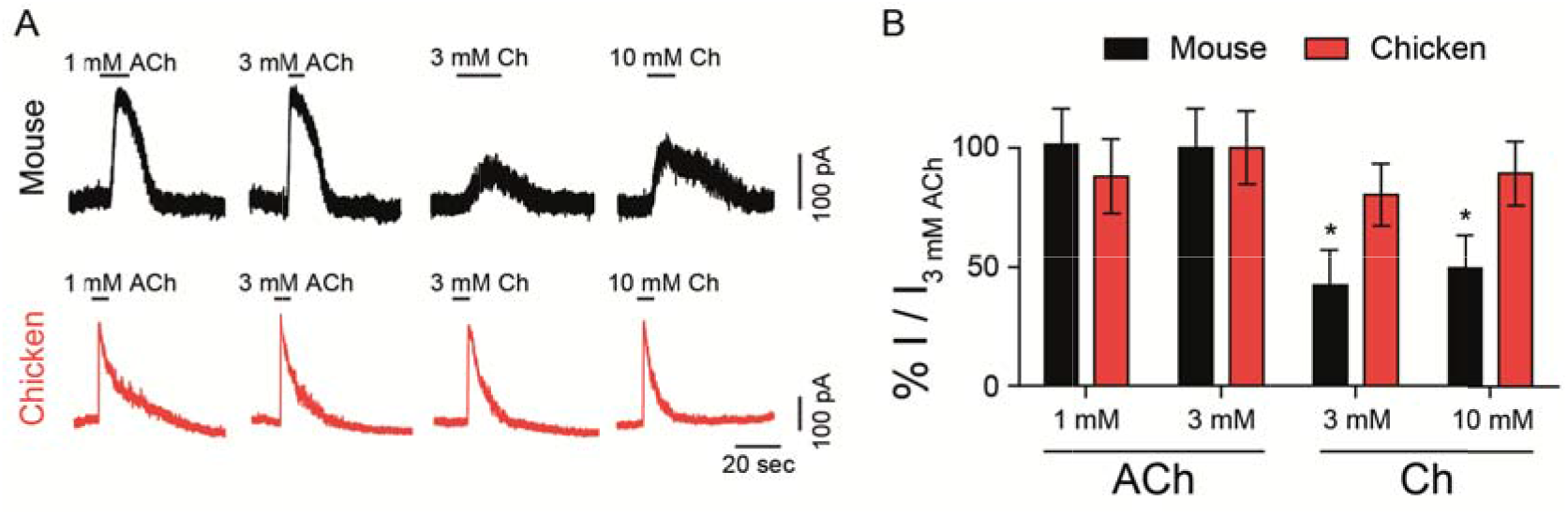
Responses of hair cells to choline. A. Representative traces of responses to ACh or choline (Ch) in mouse outer (black, top panel) and chicken short (red, bottom panel) hair cells at a holding potential of −40 mV. B. Percentage of maximum response evoked by ACh or choline normalized to the maximal peak response to 3 mM ACh for mouse and chicken hair cells. Values are mean ± S.E.M. of 6 (rat) and 4 (chicken) experiments per group. (*) Friedman’s test, vs 3 mM ACh for each species (p < 0.05).

### The α10 subunit determines choline potency and efficacy in α9α10 nAChRs

In order to determine if the agonistic characteristics of choline could be traced back to the constituent subunits of the α9α10 heteromeric receptors, we first tested the potency of choline on homomeric α9 and α10 nAChRs. Of note, mammalian α10 receptors do not assemble into functional homomeric receptors (Elgoyhen et al., 2001; Sgard et al., 2002) and were therefore not tested.

In contrast to that observed for rat heteromeric α9α10 receptors (Fig. 1D), choline behaved as a nearly full agonist of rat α9 receptors (Fig. 3A and Table 1). Thus, although less potent than ACh (EC_50_ choline 188 ± 40, n=3; ACh 12.9 ± 1.1, n=5; p=0.04 Mann-Whitney test), the maximal choline response was 87 ± 6 % (n=4, p=0.002 Kruskal-Wallis test) of that elicited by 1 mM ACh. Additionally, the EC_50_ for choline observed for the rat α9 receptor was smaller than that of the rat α9α10 receptor (Fig 1D, p=0.02 Kruskal-Wallis test). It is worth noting that choline was previously shown to be nearly a full agonist of rat α9 homomeric receptors, albeit with a higher potency than that observed here (Verbitsky et al., 2000). Differences might derive from the fact that experiments in the present work were performed in the presence of BAPTA-AM to isolate the cholinergic responses from the secondary activation of the Ca^2+^ sensitive chloride current present in oocytes (Miledi and Parker, 1984), while this was not the case in the previous work (Verbitsky et al., 2000).

**Figure 3.**
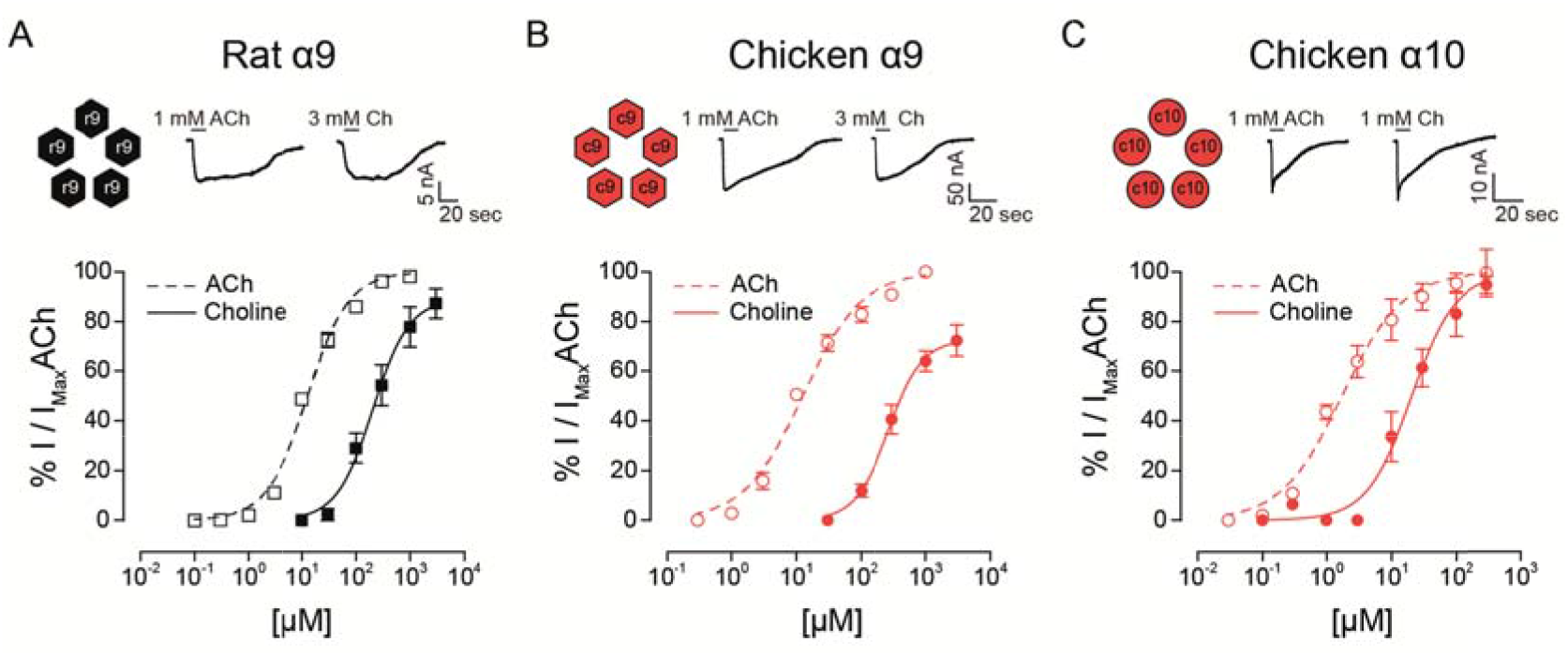
Responses of rat and chicken homomeric α9 or α10 nAChRs to choline. Top panels: representative maximal responses to ACh and choline for rat α9 (A), chicken α9 (B) and chicken α10 (C) homomeric receptors. Bottom panels: concentration-response curves for ACh (dotted lines) and choline (solid lines) for rat α9 (A), chicken α9 (B) and chicken α10 (C) homomeric receptors. Values are normalized to the maximal response to ACh obtained in each oocyte. Values are mean ± S.E.M. for 4-6 oocytes.

Choline had high efficacy in both chicken α9 and α10 homomeric receptors, similar to that observed for the chicken α9α10 heteromeric receptor (Table 1). The maximal choline response in chicken α9 receptors was 73.7 ± 5.9 % (n=6) of that obtained for 1 mM ACh (Fig. 3B), and it matched the maximal ACh response in chicken α10 receptors (99.4 ± 9.5 % (n=4); Fig. 3C). Choline potency was lower compared to that of ACh for both chicken homomeric receptors and concentration-response curves to choline were shifted to the right with higher EC_50_ values (Fig. 3B-C). For chicken α9, choline showed an EC_50_ = 280 ± 20 μM (n=6) compared to 12.9 ± 1.4 μM for ACh (n=5, p=0.004 Mann-Whitney test). In chicken α10 receptors the EC_50_ for choline was 23.2 ± 8.3 μM (n=4), compared to 2.2 ± 0.7 μM for ACh (n=5, p=0.016 Mann-Whitney test).

In summary, choline behaved as an agonist with high efficacy for homomeric receptors composed of rat α9, chicken α9 or chicken α10 subunits. This agonistic behavior was similar to that observed for chicken α9α10 receptors but differed from that of rat α9α10 heteromeric receptors.

The observation that choline behaved as a partial agonist of heteromeric rat α9α10 receptors, but a full agonist of rat α9 homomeric nAChRs, suggests that the rat α10 subunit may contain determinants responsible for the lower efficacy of choline on heteromeric α9α10 receptors. In order to test this hypothesis, we expressed rat-chicken hybrid receptors in *X. laevis* oocytes (Fig. 4). We previously determined that responses in these hybrid α9α10 receptors are indeed the result of heteromeric assemblies and not individual homomeric receptors (Lipovsek et al., 2014). As shown in Figure 4A, rat α9 – chicken α10 hybrid receptors responded to ACh with an EC_50_ of 11.6 ± 2.9 μM (n=7), similar to that of rat (EC_50_ 17.1 ± 3.1 μM, n=5, p=0.19 Kruskal-Wallis test) and chicken (EC_50_ 13.1 ± 2.6 μM, n=10, p=0.67 Kruskal-Wallis test) α9α10 nAChRs (Table 1). Distinct from that described for rat α9α10 receptors (Fig. 1D), choline produced near maximal responses in rat α9 – chicken α10 hybrid receptors (86 ± 2 % of responses to 1 mM ACh, n=6), that were similar to those observed in chicken α9α10 receptors (88 ± 7 %, n = 6). On the other hand, hybrid chicken α9 – rat α10 receptors exhibited similar concentration-response curves to ACh when compared to rat and chicken α9α10 receptors, with an EC_50_ of 18.2 ± 4.8 μM (n=4, p=0.99 and p=0.29, respectively, Kruskal-Wallis test; Fig. 4B and Table 1). However, the maximal response to choline of chicken α9 – rat α10 hybrid receptors (51 ± 6 %, n=5) was smaller than that observed for chicken α9α10 receptors (88 ± 7 %, n=10, p=0.02 Kruskal-Wallis test; Fig. 1C), with a lower efficacy, similar to that of rat α9α10 receptors (37.5 ± 2.5 %, n=9, p=0.50 Kruskal-Wallis test).

**Figure 4.**
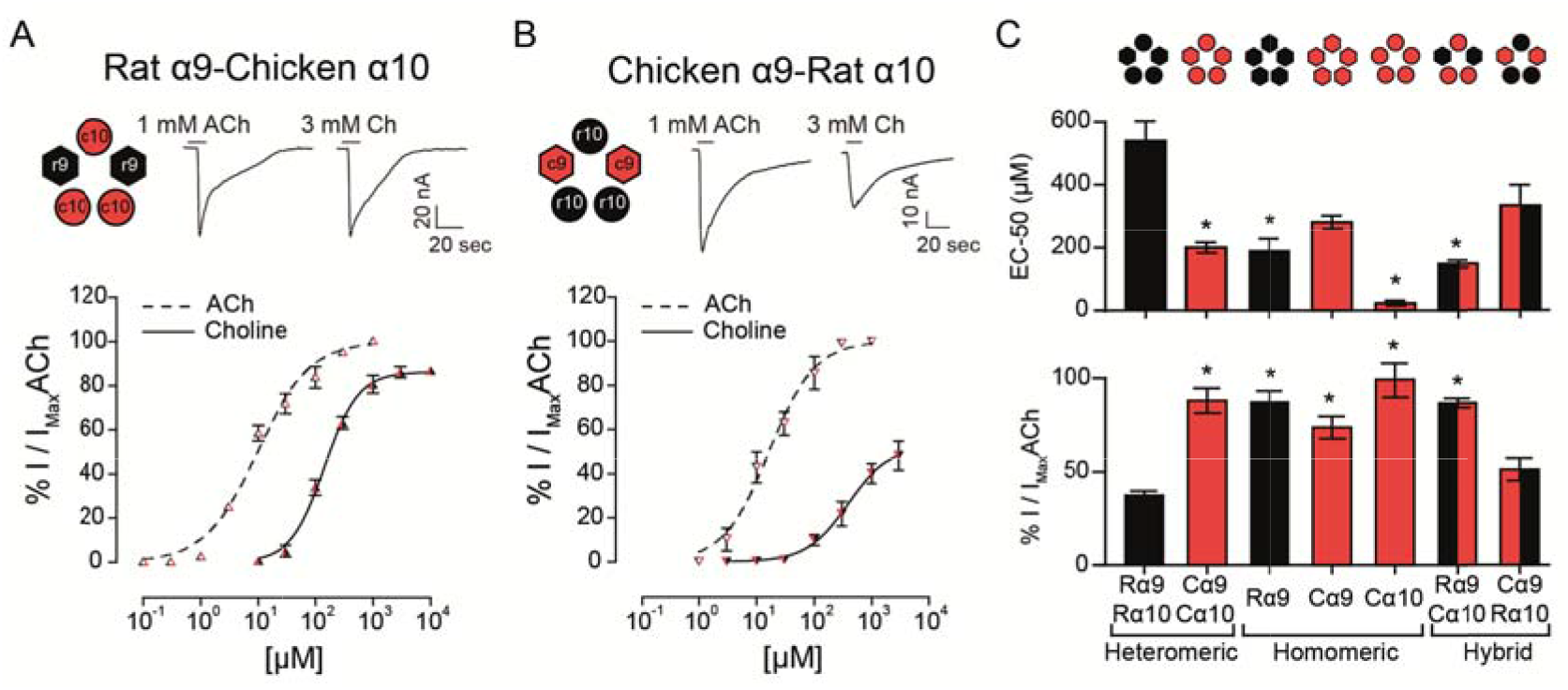
Responses of rat-chicken hybrid receptors to choline. A-B. Top panels: representative maximal responses evoked by saturating concentrations of ACh and choline in rat α9 - chicken α10 (A) and chicken α9 - rat α10 (B) hybrid receptors. Bottom panels: concentration-response curves for ACh (dotted lines) and choline (solid lines). Values are normalized to the maximal response to ACh in each oocyte. Values are mean ± S.E.M. of 4-6 oocytes per group. C. EC_50_ values for choline (Top panel) and percentage maximal choline response, normalized to the maximal response elicited by ACh (Bottom panel) for rat and chicken heteromeric α9α10 receptors, rat α9, chicken α9 and chicken α10 homomeric receptors, and rat α9 – chicken α10 and chicken α9 – rat α10 hybrid receptors. (*) Kruskal-Wallis test, vs rat α9α10 receptor (p<0.05).

In summary, both the potency and efficacy of choline were lowest for rat heteromeric α9α10 receptors than all other receptors, with the exception of heteromeric hybrid receptors containing rat α10 subunits, therefore suggesting that the rat α10 subunit may hold the molecular determinants of choline’s partial agonism (Fig. 4C).

### Lower frequency of choline binding at orthosteric sites holding rat α10 subunits as complementary components

To gain further insight into the mechanism underlying the drop in choline agonism on receptors containing the rat α10 subunit, we performed molecular docking simulations and analyzed the interaction of ACh and choline with the orthosteric binding site. To this end, we used homology models of the extracellular domains of rat and chicken α9α10 receptors with subunit arrangements corresponding to the four possible binding site interfaces (principal/complementary components): α9/α9, α9/α10, α10/α9 and α10/α10 as previously reported (Boffi et al., 2017). We then performed molecular docking analysis of the interaction of ACh or choline with the homology-modeled subunit interfaces to evaluate the orientation, the best binding energy (BBE) and the frequency of conformations that bind the agonist in a favorable orientation within the binding pocket. The conformations considered as favorable were those that showed typical cation-π interactions between the amino group of the ligands and aromatic residues of the binding pocket (W55, from the complementary face, and Y93, W149, Y190 and/or Y197 from the principal face) required for ACh responses (Dougherty, 2007; Olsen et al., 2014).

As reported previously (Boffi et al., 2017), both for rat and chicken receptors, ACh docking resulted in an energetically favorable model with ACh oriented with its quaternary amine toward the membrane side or lower part of the cleft and the negative charge oriented upwards in the cleft (Fig. 5A and B), similar to ACh in AChBP (Olsen et al., 2014). In this orientation, ACh showed BBE between −3.5 and −6 kcal/mol for the different binding interfaces (Fig. 5E) and its positively charged group showed the potential to form the typical cation-π interactions with the main aromatic residues Y93, W149, Y190, Y197 and W55 (Fig. 5B and C). Thus, as reported for other nAChRs, the orientation of ACh in the binding pocket of rat and chicken α9α10 receptors is favorable for efficacious activation (Fig. 5 and (Boffi et al., 2017)).

**Figure 5.**
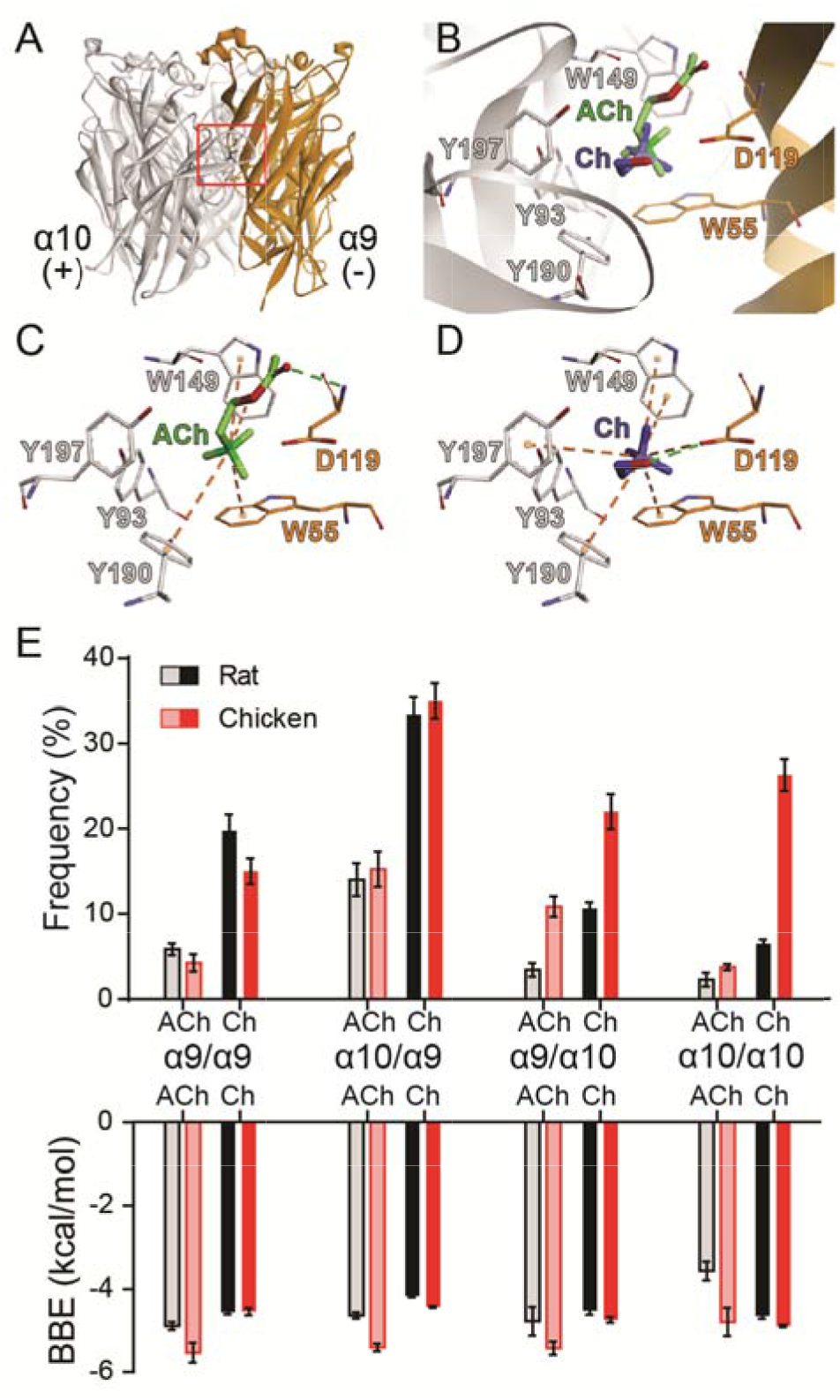
Docking of ACh and choline into the binding sites of rat and chicken a9a10 receptors. A. Ribbon structure of the extracellular domain of rat α10 and α9 subunits showing an interface formed by the principal (+) component of the α10 subunit and the complementary (−) component of the α9 subunit. The location of the orthosteric binding site with a bound ACh is highlighted by the red box. B. Detailed view of the binding domain highlighted by the red square in A. ACh (green) and the most frequent orientation of choline (blue) are shown docked into the binding pocket. C-D. Molecular interactions of ACh (C) and choline (D) with residues in the binding site. Cation-π interactions are illustrated by orange dashed lines and H-bonds by green dashed lines. E. Frequency (as percentage) of the most frequent orientation (Top panel) and best binding energy (BBE; Bottom panel) obtained for three separate simulations for ACh and choline docking into each of the binding interfaces, comprised of (principal/complementary components) α9/α9, α10/α9, α9/α10 and α10/α10 rat (black) or chicken (red) subunits. Values are mean ± SEM of at least three different docking simulations for each interface.

For all interfaces, choline docked into the orthosteric binding site with two different orientations. The least frequent orientation was similar to that described for choline docked into the muscle nAChR (Bruhova and Auerbach, 2017). In the most frequent and lowest BBE conformation, choline was located more horizontally with respect to the membrane compared to ACh, oriented towards the C-loop containing Y190 and Y197, with the quaternary ammonium placed in the aromatic cavity (Fig. 5B and D). This orientation has also been described for other ligands in different nAChR models (Tomaselli et al., 1991; Lester et al., 2004; Hernando et al., 2012). In this orientation, choline showed the potential to make typical cation-π interactions with the aromatic residues at the principal (W149, Y93, Y190, Y197) and complementary (W55) components (Fig. 5D). Whereas ACh formed an H-bond with the backbone amine group of residue D119 (Boffi, 2017 and Fig. 5B and C), choline formed an H-bond with the carboxyl group of D119 (Fig. 5B and D). Altogether, the orientation and potential interactions with residues indicate that choline adopts a favorable conformation (though different to that of ACh) for eliciting receptor activation, with lower BBEs than those observed for ACh (between −4 and −4.5 kcal/ mol for all interfaces).

The orientation and main interactions of choline with residues at the binding site were identical for all interfaces and between chicken and rat receptors. Also, while the BBE for choline did not show significant inter-species differences across the alternate interfaces, we observed variations in the frequency at which choline bound with favorable conformations to the different interfaces (Fig. 5E). Those binding interfaces containing α9 as the complementary subunit (α9/α9 and α10/α9) showed high frequency of choline binding in the favorable conformations both for chicken and rat subunits (Fig. 5E). In contrast, differences were encountered at interfaces in which α10 provided the complementary side (α9/α10 and α10/α10). While each agonist binding frequency for chicken interfaces was similar to that observed with α9 as the complementary subunit, it was significantly lower for interfaces showing a rat α10 complementary subunit (Fig. 5F).

Thus, our *in silico* studies focused on the binding site interfaces suggest that the differences of choline responses may be governed by the distinct choline orientation and the differential contribution of binding interfaces containing α10 as the complementary subunit. While in chicken α9α10 receptors a high frequency of choline binding was observed for all conformations of binding sites, in rat α9α10 receptors the frequency of choline binding was significantly lower at α9/α10 and α10/α10 sites, suggesting that the lower choline agonism observed in rat α10 containing receptors (Figures 3 and 4) may be related to a reduced capability of the rat α10 subunit to efficiently operate as the complementary side during agonist binding and receptor gating. Consequently, a reduction in the binding frequency of agonists at specific interfaces might determine differential responses in the context of receptor function.

### The extracellular region of the rat α10 subunit underpins lower choline efficacy

The N-terminal and TM2-TM3 extracellular domains of nicotinic receptors bare residues involved both in agonist binding and coupling to pore opening (Karlin, 2002; Bouzat et al., 2004), and most likely contain determinants for agonist efficacy (Gupta et al., 2017; Mukhtasimova and Sine, 2018). Given that choline partial agonism was observed when rat α10 subunits are present in the heteromeric receptors (Fig. 4C) and that, although it binds to all binding site combinations, the frequency of favorable choline binding was lower at interfaces in which rat α10 provided the complementary face (Fig. 5F), we hypothesized that a receptor in which no binding site contains α10 components would show strong choline agonism. To test this, we engineered a rat α9-α10 (α10_x_) chimeric subunit in which the entire N-terminal and TM2-TM3 extracellular regions of the rat α10 subunit were replaced by the corresponding domains of the rat α9 subunit (Fig. 6A). The chimeric subunit did not form functional homomeric receptors by itself, nor did it do so when co-expressed with the rat α10 subunit. However, when co-expressed with the rat α9 subunit, strong ACh responses were recorded for rat α9α10_x_ receptors (Fig. 6B), with I_max_ = 269 ± 103 nA (n=5). This contrasts the maximal ACh responses observed for homomeric rat α9 receptors (I_max_ of 14 ± 2 nA, n=4 and see (Elgoyhen et al., 2001), indicating that, when co-expressed, rat α9 and chimeric α10_x_ subunits form heteromeric assemblies. Concentration-response curves for ACh on rat α9α10_x_ receptors were similar to those of rat α9α10 wild-type receptors with an EC_50_ of 15.6 ± 6.5 μM (n=4, p=0.50 Kruskal-Wallis test) (Fig. 6C). Interestingly, responses to choline of the rat α9α10_x_ chimeric receptor resembled those exhibited by rat α9 homomeric receptors and differed from those of rat α9α10 receptors. Thus, the EC_50_ for choline on α9α10_x_ receptors was 160.2 ± 7.6 μM (n=5), significantly smaller than that in rat α9α10 receptors (541 ± 62 μM, n=10, p=0.007, Kruskal-Wallis test) and similar to that observed in homomeric rat α9 receptors (Fig. 6C and Table 1). Likewise, choline-evoked maximal response of 74 ± 1 % (n=5) in α9α10_x_ receptors was higher than that of rat α9α10 (38 ± 3 %, n=9, p=0.04 Kruskal-Wallis test) but similar to that observed in rat α9 homomeric receptors (87 ± 6 %, n=4, p=0.27, Kruskal-Wallis test) (Fig. 6C and Table 1). Taken together, the experimental evaluation of ACh and choline agonism in the chimeric receptor indicates that the extracellular region of the rat α10 subunit bears determinants that reduce the efficacy of choline on the α9α10 receptor.

**Figure 6.**
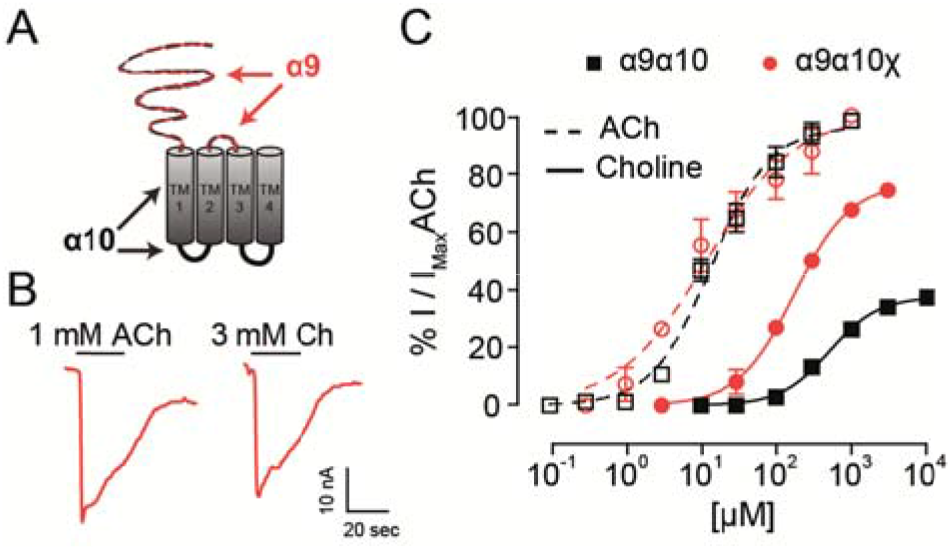
Responses of rat α9α10_x_ chimeric receptor to choline. A. Diagram of an α9α10_x_ chimeric subunit showing the regions corresponding to the α9 subunit sequence (red) and α10 subunit sequence (black). B. Representative maximal responses to ACh (left) and choline (right) for the α9α10_x_ chimeric receptor. C. Concentration response-curves for ACh (dotted lines) and choline (solid lines) for α9α10_x_ chimeric receptors (red circles) and wild-type α9α10 receptors (black squares). Values were normalized to the maximal response to ACh in each oocyte. Values are mean ± S.E.M. of 5-9 oocytes per group.

### The α10 complementary component participates in activation by choline of rat α9α10 nAChRs

In a previous work we showed that rat α10 subunits do not contribute with complementary faces to the ACh binding site (Boffi et al., 2017). In contrast, in the present study we have so far shown that the extracellular domain of the rat α10 subunit is responsible for the decrease in choline agonism in rat α9α10 receptors. Moreover, *in silico* docking showed that the frequency of choline binding in the correct conformation is lower at interfaces where the complementary component is provided by rat α10 subunits (Fig. 5E), suggesting that the lower choline agonism observed in rat α9α10 receptors may result from sub-functional α9/α10 or α10/α10 binding sites. Therefore, to evaluate experimentally choline’s requirement for functional binding sites containing α10 complementary components, we studied responses to choline on rat α9α10 heteromeric receptors holding the W55T mutation in the rat α10 subunits (Fig. 7A). W55 is located within loop D of the complementary component of the binding site, which by directly interacting with the agonist is crucial for binding-gating transduction (Olsen et al., 2014). While responses to ACh showed no significant changes in the α9α10W55T receptor compared to wild type α9α10 receptors (Boffi et al., 2017), Fig. 7A and B and Table 1 - EC_50_ α9α10: 17.1 ± 3.1 μM, n=5; EC_50_ α9α10W55T: 37.8 ± 6.2 μM, n=8; p=0.06, Kruskal-Wallis test), responses to choline were nearly abolished in mutant receptors. Thus, responses to 10 mM choline were 7 ± 2 % (n=5) of those elicited by 1 mM ACh (Fig. 7B). This indicates that the contribution of the complementary face of the rat α10 subunit to the agonistic effect of ACh and choline is non-equivalent. Thus, whereas rat α10 complementary faces are not required for ACh activation of rat α9α10 receptors (see also (Boffi et al., 2017), intact rat α10 complementary components are required for choline agonistic effect.

**Figure 7.**
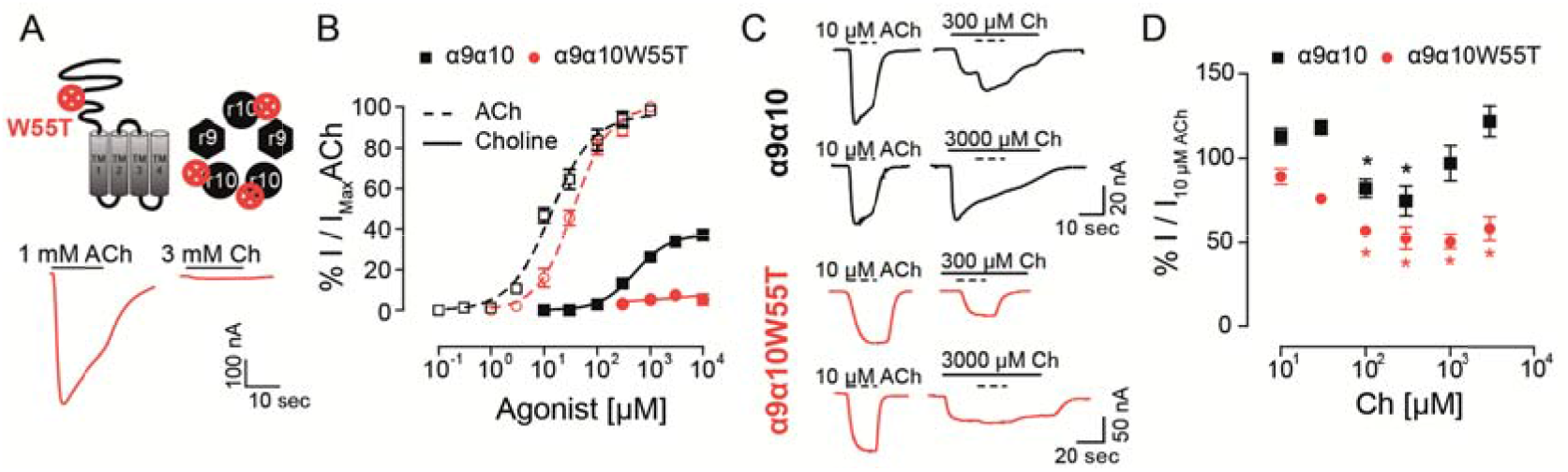
Responses of the rat α9α10W55T receptor to choline. A. Diagram of an α10 subunit illustrating the position of the W55T mutation, localized to the complementary component of the agonist binding site within the pentameric assembly (Top panel). Representative maximal responses to ACh (left) and choline (right) for the α9α10W55T receptor (Bottom panel). B. Concentration response-curves to ACh (dotted lines) and choline (solid lines) for the α9α10W55T receptor (red circles) and wild-type α9α10 receptors (black squares). Values were normalized to the maximal response to ACh in each oocyte. Values are mean ± S.E.M. of 5-9 oocytes per group. C. Representative responses to 10 μM ACh (dotted lines) alone or on top of responses to 300 or 3000 μM choline in oocytes expressing wild-type α9α10 or α9α10W55T receptors. D. Normalized responses to 10 μM ACh obtained in the presence of increasing concentrations of choline for wild-type α9α10 and α9α10W55T receptors. Responses were normalized to responses to10 μM ACh in the absence of choline for each oocyte.

Finally, to evaluate whether choline interacts with the α10/α9 intact interfaces in the α9α10W55T receptor, and therefore also requires them to elicit its reduced agonism, we analyzed responses to 10 μM ACh in the presence of increasing concentrations of choline for both rat α9α10 and α9α10W55T receptors (Fig. 7C and D). ACh was added once a maximal response to choline was observed. The maximal response obtained during the co-application was normalized to the response to 10 μM ACh (Fig. 7C). At low choline concentrations, that do not trigger channel opening (< 100 μM), ACh responses became increasingly smaller as choline concentration increased, both in the α9α10 and the α9α10W55T receptors, suggesting that choline is capable of competing with ACh for the available binding sites. However, in the presence of higher choline concentrations (>300 μM), total responses in α9α10 receptors increased while they remained unchanged for the mutant receptor (Fig. 7D). The V-shaped effect observed in the α9α10 receptor, where an EC_50_ ACh response is blocked at low concentrations of choline and summation of both agonist responses occurs at higher choline concentrations, is consistent with choline acting as a partial agonist (Zhu, 2005) and suggests that the increase in combined response may be due to choline binding to the α9/α10 and α10/α10 sites that are spared by ACh. Conversely, the lack of increase of ACh responses at higher choline concentrations in the α9α10W55T mutant receptor indicates a loss of choline agonism that may result from the unavailability of additional α9/α10 and α10/α10 sites due to the W55T mutation.

Overall, the differences observed in the responses to ACh and/or choline between α9α10 wild type and α9α10W55T mutant receptors indicate that the complementary faces of α10 subunits are required for the partial agonistic effect of choline on α9α10 receptors, therefore suggesting that choline necessitates higher binding site occupancy compared to ACh to elicit channel gating of rat α9α10 nAChRs.

## Discussion

The present work shows that the mammalian (rat) α10 nAChR subunit is responsible for a reduced efficacy of choline when assembled into heteromeric α9α10 receptors. Thus, whereas choline is a full agonist of rat α9 homomeric receptors, it is a partial agonist of rat α9α10 nAChRs. This is further supported by hybrid receptors, on which choline is a full agonist of rat α9 – chicken α10 receptors and the efficacy in chicken heteromeric receptors is reduced when the chicken α10 subunit is replaced by the rat α10 subunit in the chicken α9 – rat α10 assembly. The reduced agonist efficacy determined by the rat α10 subunit is probably extended to other compounds that behave as partial agonists, as evidenced in the present work for DMPP. These results add to the differential functional properties between mammalian and non-mammalian vertebrate α9α10 nAChRs that have been shaped by the more prevalent occurrence of non-synonymous amino acid substitutions during the evolution of mammalian subunits (Franchini and Elgoyhen, 2006; Elgoyhen and Franchini, 2011; Lipovsek et al., 2012; Lipovsek et al., 2014; Marcovich et al., 2020).

Full and partial agonists evoke distinct structural changes in opening the muscle acetylcholine receptor channel (Mukhtasimova and Sine, 2018). This is further supported by the observation that the αE45R mutation within the binding–gating transduction domain of this receptor attenuates channel opening by a full agonist, whereas it enhances channel opening by a partial agonist (Mukhtasimova and Sine, 2013). It was therefore suggested that, due to differences in size and/or chemical structure, different agonists of the same receptor might bind in different conformations and/or strength to the ligand interaction pocket (Mukhtasimova and Sine, 2018). This resembles our observations on the effect of mutating W55 in the rat α10 subunit which greatly impairs the response of the α9α10 receptor to choline but not to ACh.

This functional non-equivalence between agonists is further supported by docking analysis, which shows that ACh and choline adopt different conformations at the binding site of α9α10 receptors. This result fully agrees with previous studies in the muscle nAChR (Bruhova and Auerbach, 2017), for which choline and ACh sit at different orientations at the binding site. In our model, choline had two probable alternate orientations compared to ACh with the least frequent one resembling the one described for the muscle nAChR (Bruhova and Auerbach, 2017). Furthermore, the non-equivalence of the agonist binding sites for ACh and choline is reflected in differential atomic interactions. Thus, similar to ACh, choline can make interactions at the aromatic cage. However, whereas ACh makes a H-bond with the backbone amine group of D119 of the complementary face, choline shows the potential to make a H-bond with the carboxyl group of the same residue (Fig. 5B and D). Similar to that reported for the muscle ACh receptor (Bruhova and Auerbach, 2017), this different orientation leads to the displacement of the quaternary ammonium of choline away from a favorable position in the aromatic cage compared to ACh. This might lead to a weaker interaction with all aromatic rings of the binding site, accounting for the lower apparent affinity of choline compared to ACh for the α9α10 receptor. Moreover, due to the particular orientation of choline towards the C-loop when compared to ACh, differential interactions with residues of the complementary face and/or degree of closure of the C-loop required for channel gating (Thompson et al., 2010), might be involved in the reduced choline efficacy.

Our combined *in silico* and experimental observations allow us further insight into the differential responses to ACh and choline of rat α9α10 nAChRs. Whereas the complementary face of the α10 subunit does not play an important role in the activation of the receptor by ACh, it is strictly required for choline responses, as shown by the results of the W55T mutation. ACh adopts a different and more favorable conformation, better placed within the aromatic pocket compared to choline, and would therefore only require occupying two of the five binding sites, sparing binding sites in which the complementary face is provided by the α10 subunit. However, since choline adopts a less favorable conformation, it probably requires higher binding site occupancy for full efficacy. Differences in agonist efficacy according to available binding sites have been reported for other nAChRs. For example, the α4β2 nAChR in the (α4)_3_(β2)_2_ stoichiometry contains three functional agonist binding sites for ACh (Harpsøe et al., 2011; Mazzaferro et al., 2011), and the engagement of all three agonist sites produces maximal activation. The agonist site at the α4/α4 interface is a key determinant of agonist efficacy as occupancy of this site increases agonist efficacy, whereas exclusion from the site leads to partial agonism (Mazzaferro et al., 2014). Another example is provided by the α7 receptor. Even though maximal activation requires occupancy of three binding sites in the α7-5HT_3_A chimera (Rayes et al., 2009), only one is required in the α7 nAChR (Andersen et al., 2013). Thus, the relationship between binding site occupancy and maximal response differs between nAChRs. Moreover, and in light of our results, we propose that the degree of occupancy required for maximal responses varies with the type of agonist.

Hair cell α9α10 nAChRs are distinct from other nicotinic receptors in that a greater divergence in their coding sequence has translated into differential functional properties across clades (Lipovsek et al., 2012; Lipovsek et al., 2014; Marcovich et al., 2020). Our observations here that choline, the degradation product of ACh, has differential agonistic effects on rodent vs avian α9α10 receptors, suggests different scenarios for the workings of the respective efferent synapses. Recordings on native cholinergic responses recapitulated our observations in recombinant rat and chick α9α10 receptors. While choline elicited maximal responses in chicken hair cells, it behaved as a partial agonist of the native nicotinic receptors present in hair cells of mice (Fig. 2). These results imply that in chicken efferent synapses to (mostly) short hair cells, the release of ACh from efferent terminals triggers α9α10 receptors that would continue to be activated in response to the choline produced by ACh degradation due to acetylcholinesterase (AChE) activity, until it is removed from the synaptic cleft. This would result in longer post-synaptic responses subjected to large variations and poor temporal tuning (Katz and Miledi, 1973). In contrast, in mammalian efferent olivocochlear synapses, the degradation of ACh to choline would limit the time-course and improve the reliability of the cholinergic response. Given that choline is not able to fully activate the rodent α9α10 receptor, and in the presence of ACh acts as a competitive antagonist, the termination of α9α10 responses would therefore be dictated by the fast kinetics of AChE activity (Hall, 1973). An apical to basal concentration and isoform diversity gradient of AChE has been described in the mouse cochlea (Emmerling and Sobkowicz, 1988). Notably, the high-frequency basal region exhibits higher concentrations of this enzyme and is enriched in isoforms with faster kinetics, underscoring the relevance of fine temporal tuning of efferent modulation for high frequency sound detection (Emmerling and Sobkowicz, 1988). Additionally, BK channels, that display larger currents with faster kinetics than SK2 channels, participate in efferent synaptic inhibition in higher frequency regions of the cochlea, supporting the notion that an accurate control of OHCs membrane potential is required for the amplification and modulation of high frequency hearing (Rohmann et al., 2015). Finally, compared to chicken, the mammalian α9α10 receptor shows greater desensitization to ACh and higher calcium permeability (Lipovsek et al., 2012; Lipovsek et al., 2014; Marcovich et al., 2020). In addition, the postsynaptic space delimited by a closely juxtaposed subsynaptic cistern is more restricted in mammals (Fuchs et al., 2014; Im et al., 2014), which may contribute to elicit highly localized α9α10-dependent rises in calcium concentration uncoupled from internal calcium stores (Moglie et al., 2018; Moglie et al., 2020). Together with the poorer choline agonism, these multiple functional adaptations may have contributed to “tighter” ACh responses, which may prove fundamental to faithfully reproduce the high frequency activity of efferent medial olivocochlear fibers (Ballestero et al., 2011), therefore fine-tuning the modulation of the OHC cochlear amplifier and contributing to the expansion of the mammalian hearing range.

The lower agonistic action of choline on mammalian α9α10 receptors is likely due to the accumulation of amino acid changes within the mammalian α10 subunit (Franchini and Elgoyhen, 2006; Lipovsek et al., 2012) that rendered a subfunctional contribution of α10 as a complementary component subunit. Our *in silico* and experimental analysis of choline agonism indicate that, for it to trigger a full response, it needs to bind to α9/α10 and/or α10/α10 sites, therefore requiring sites holding α10 complementary interfaces. This is in contrast with the agonism by ACh, that only requires binding to α10/α9 (or α9/α9) sites formed by an α10 (or α9) principal component and α9 complementary component (Boffi et al., 2017). We hypothesize that amino acid changes within the α10 subunit that affect agonist binding and/or receptor triggering from the additional binding sites (i.e. those that have an α10 subunit as a complementary component) would not be deleterious and therefore not under negative selection, as they would not affect the main response to ACh, given that the sensitivity for this agonist is the same in the different species (Lipovsek et al., 2012; Marcovich et al., 2020). In this context, a scenario could be proposed in which strong positive selection pressure for the loss of the agonistic function of choline, a likely functional requirement for fine-tuned high-frequency efferent olivocochlear activity, may have been the driver for the accumulation of coding sequence changes within the extracellular domain of mammalian α10 subunits (Franchini and Elgoyhen, 2006). This then resulted in the loss of choline acting as a full agonist for mammalian α9α10 receptors, without affecting the triggering of ACh responses.

## Acknowledgments

We thank Paul A. Fuchs for assistance in the performance of chicken hair cell recordings. This work was supported by Agencia Nacional de Promoción Científicas y Técnicas, Argentina, the Scientific Grand Prize of the Fondation Pour l’Audition, and NIH grant R01 DC001508 (Paul Fuchs PI and ABE co-PI) to A.B.E.

## Notes

### Competing Interest Statement

The authors have declared no competing interest.

## References

Albuquerque, E.X., Pereira, E.F., Braga, M.F., and Alkondon, M. (1998). Contribution of nicotinic receptors to the function of synapses in the central nervous system: the action of choline as a selective agonist of alpha 7 receptors. J Physiol Paris 92(3-4), 309–316. doi: 10.1016/s0928-4257(98)80039-9.

Andersen, N., Corradi, J., Sine, S.M., and Bouzat, C. (2013). Stoichiometry for activation of neuronal α7 nicotinic receptors. Proc Natl Acad Sci U S A 110(51), 20819–20824. doi: 10.1073/pnas.1315775110.

Arnold, K., Bordoli, L., Kopp, J., and Schwede, T. (2006). The SWISS-MODEL workspace: a web-based environment for protein structure homology modelling. Bioinformatics 22(2), 195–201. doi: 10.1093/bioinformatics/bti770.

Ballestero, J., Zorrilla de San Martin, J., Goutman, J., Elgoyhen, A.B., Fuchs, P.A., and Katz, E. (2011). Short-term synaptic plasticity regulates the level of olivocochlear inhibition to auditory hair cells. J Neurosci 31(41), 14763–14774. doi: 10.1523/jneurosci.6788-10.2011.

Ballestero, J.A., Plazas, P.V., Kracun, S., Gomez-Casati, M.E., Taranda, J., Rothlin, C.V., et al. (2005). Effects of quinine, quinidine, and chloroquine on alpha9alpha10 nicotinic cholinergic receptors. Mol Pharmacol 68(3), 822–829. doi: mol.105.014431 [pii] 10.1124/mol.105.014431 [doi].

Boffi, J.C., Marcovich, I., Gill-Thind, J.K., Corradi, J., Collins, T., Lipovsek, M.M., et al. (2017). Differential Contribution of Subunit Interfaces to alpha9alpha10 Nicotinic Acetylcholine Receptor Function. Mol Pharmacol 91(3), 250–262. doi: 10.1124/mol.116.107482.

Bordoli, L., Kiefer, F., Arnold, K., Benkert, P., Battey, J., and Schwede, T. (2009). Protein structure homology modeling using SWISS-MODEL workspace. Nat Protoc 4(1), 1–13. doi: 10.1038/nprot.2008.197.

Bouzat, C., Gumilar, F., Spitzmaul, G., Wang, H.L., Rayes, D., Hansen, S.B., et al. (2004). Coupling of agonist binding to channel gating in an ACh-binding protein linked to an ion channel. Nature 430(7002), 896–900. doi: 10.1038/nature02753.

Bruhova, I., and Auerbach, A. (2017). Molecular recognition at cholinergic synapses: acetylcholine versus choline. J Physiol 595(4), 1253–1261. doi: 10.1113/jp273291.

Dougherty, D.A. (2007). Cation-pi interactions involving aromatic amino acids. J Nutr 137(6 Suppl 1), 1504S–1508S; discussion 1516S-1517S. doi: 10.1093/jn/137.6.1504S.

Dulon, D., and Lenoir, M. (1996). Cholinergic responses in developing outer hair cells of the rat cochlea. European J. Neurosci 8, 1945–1952.

Elgoyhen, A.B., and Franchini, L.F. (2011). Prestin and the cholinergic receptor of hair cells: Positively-selected proteins in mammals. Hear Res 273(1-2), 100–108. doi: S0378-5955(09)00328-1 [pii] 10.1016/j.heares.2009.12.028 [doi].

Elgoyhen, A.B., Johnson, D.S., Boulter, J., Vetter, D.E., and Heinemann, S. (1994). a9: an acetylcholine receptor with novel pharmacological properties expressed in rat cochlear hair cells. Cell 79, 705–715.

Elgoyhen, A.B., and Katz, E. (2012). The efferent medial olivocochlear-hair cell synapse. J Physiol Paris 106(1-2), 47–56. doi: 10.1016/j.jphysparis.2011.06.001.

Elgoyhen, A.B., Katz, E., and Fuchs, P.A. (2009). The nicotinic receptor of cochlear hair cells: a possible pharmacotherapeutic target? Biochem Pharmacol 78(7), 712–719. doi: S0006-2952(09)00436-5 [pii] 10.1016/j.bcp.2009.05.023 [doi].

Elgoyhen, A.B., Vetter, D., Katz, E., Rothlin, C., Heinemann, S., and Boulter, J. (2001). Alpha 10: A determinant of nicotinic cholinergic receptor function in mammalian vestibular and cochlear mechanosensory hair cells. Proc Natl Acad Sci USA 98, 3501–3506.

Emmerling, M.R., and Sobkowicz, H.M. (1988). Differentiation and distribution of acetylcholinesterase molecular forms in the mouse cochlea. Hear Res 32(2-3), 137–145. doi: 10.1016/0378-5955(88)90086-x.

Franchini, L.F., and Elgoyhen, A.B. (2006). Adaptive evolution in mammalian proteins involved in cochlear outer hair cell electromotility. Mol Phylogenet Evol 41(3), 622–635. doi: S1055-7903(06)00201-6 [pii] 10.1016/j.ympev.2006.05.042 [doi].

Fuchs, P.A., Lehar, M., and Hiel, H. (2014). Ultrastructure of cisternal synapses on outer hair cells of the mouse cochlea. J Comp Neurol 522(3), 717–729. doi: 10.1002/cne.23478.

Fuchs, P.A., and Murrow, B.W. (1992). A novel cholinergic receptor mediates inhibition of chick cochlear hair cells. Proc. R. Soc. Lond. B 248, 35–40.

Gomez-Casati, M.E., Fuchs, P.A., Elgoyhen, A.B., and Katz, E. (2005). Biophysical and pharmacological characterization of nicotinic cholinergic receptors in cochlear inner hair cells. J Physiol 566, 103–118.

Goutman, J.D., Elgoyhen, A.B., and Gomez-Casati, M.E. (2015). Cochlear hair cells: The sound-sensing machines. FEBS Lett 589(22), 3354–3361. doi: 10.1016/j.febslet.2015.08.030.

Guex, N., and Peitsch, M.C. (1997). SWISS-MODEL and the Swiss-PdbViewer: an environment for comparative protein modeling. Electrophoresis 18(15), 2714–2723. doi: 10.1002/elps.1150181505.

Gupta, S., Chakraborty, S., Vij, R., and Auerbach, A. (2017). A mechanism for acetylcholine receptor gating based on structure, coupling, phi, and flip. J Gen Physiol 149(1), 85–103. doi: 10.1085/jgp.201611673.

Hall, Z.W. (1973). Multiple forms of acetylcholinesterase and their distribution in endplate and non-endplate regions of rat diaphragm muscle. J Neurobiol 4(4), 343–361. doi: 10.1002/neu.480040404.

Harpsøe, K., Ahring, P.K., Christensen, J.K., Jensen, M.L., Peters, D., and Balle, T. (2011). Unraveling the high- and low-sensitivity agonist responses of nicotinic acetylcholine receptors. J Neurosci 31(30), 10759–10766. doi: 10.1523/jneurosci.1509-11.2011.

Hernando, G., Bergé, I., Rayes, D., and Bouzat, C. (2012). Contribution of subunits to Caenorhabditis elegans levamisole-sensitive nicotinic receptor function. Mol Pharmacol 82(3), 550–560. doi: 10.1124/mol.112.079962.

Horton, R.M., Hunt, H.D., Ho, S.N., Pullen, J.K., and Pease, L.R. (1989). Engineering hybrid genes without the use of restriction enzymes: gene splicing by overlap extension. Gene (Amst.) 77, 61–68.

Humphrey, W., Dalke, A., and Schulten, K. (1996). VMD: visual molecular dynamics. J Mol Graph 14(1), 33–38, 27-38. doi: 10.1016/0263-7855(96)00018-5.

Im, G.J., Moskowitz, H.S., Lehar, M., Hiel, H., and Fuchs, P.A. (2014). Synaptic calcium regulation in hair cells of the chicken basilar papilla. J Neurosci 34(50), 16688–16697. doi: 10.1523/jneurosci.2615-14.2014.

Karlin, A. (2002). Ion channel structure: emerging structure of the nicotinic acetylcholine receptors. Nature Reviews Neurosc 3, 102–114.

Katz, B., and Miledi, R. (1973). The binding of acetylcholine to receptors and its removal from the synaptic cleft. J Physiol 231(3), 549–574. doi: 10.1113/jphysiol.1973.sp010248.

Katz, E., and Elgoyhen, A.B. (2014). Short-term plasticity and modulation of synaptic transmission at mammalian inhibitory cholinergic olivocochlear synapses. Front Syst Neurosci 8, 224. doi: 10.3389/fnsys.2014.00224.

Katz, E., Elgoyhen, A.B., Gomez-Casati, M.E., Knipper, M., Vetter, D.E., Fuchs, P.A., et al. (2004). Developmental regulation of nicotinic synapses on cochlear inner hair cells. J Neurosci 24(36), 7814–7820.

Lester, H.A., Dibas, M.I., Dahan, D.S., Leite, J.F., and Dougherty, D.A. (2004). Cys-loop receptors: new twists and turns. Trends Neurosci 27(6), 329–336. doi: 10.1016/j.tins.2004.04.002.

Lipovsek, M., Fierro, A., Perez, E.G., Boffi, J.C., Millar, N.S., Fuchs, P.A., et al. (2014). Tracking the molecular evolution of calcium permeability in a nicotinic acetylcholine receptor. Mol Biol Evol 31(12), 3250–3265. doi: 10.1093/molbev/msu258.

Lipovsek, M., Im, G.J., Franchini, L.F., Pisciottano, F., Katz, E., Fuchs, P.A., et al. (2012). Phylogenetic differences in calcium permeability of the auditory hair cell cholinergic nicotinic receptor. Proc Natl Acad Sci U S A 109(11), 4308–4313. doi: 10.1073/pnas.1115488109.

Manley, G.A. (2000). Cochlear mechanisms from a phylogenetic viewpoint. Proc Natl Acad Sci U S A 97(22), 11736–11743.

Marcovich, I., Moglie, M.J., Carpaneto Freixas, A.E., Trigila, A.P., Franchini, L.F., Plazas, P.V., et al. (2020). Distinct Evolutionary Trajectories of Neuronal and Hair Cell Nicotinic Acetylcholine Receptors. Mol Biol Evol 37(4), 1070–1089. doi: 10.1093/molbev/msz290.

Mazzaferro, S., Benallegue, N., Carbone, A., Gasparri, F., Vijayan, R., Biggin, P.C., et al. (2011). Additional acetylcholine (ACh) binding site at alpha4/alpha4 interface of (alpha4beta2)2alpha4 nicotinic receptor influences agonist sensitivity. J Biol Chem 286(35), 31043–31054. doi: 10.1074/jbc.M111.262014.

Mazzaferro, S., Gasparri, F., New, K., Alcaino, C., Faundez, M., Iturriaga Vasquez, P., et al. (2014). Non-equivalent ligand selectivity of agonist sites in (α4β2)2α4 nicotinic acetylcholine receptors: a key determinant of agonist efficacy. J Biol Chem 289(31), 21795–21806. doi: 10.1074/jbc.M114.555136.

Miledi, R., and Parker, I. (1984). Chloride current induced by injection of calcium into Xenopus oocytes. J. Physiol. (Lond). 357, 173–183.

Moglie, M.J., Fuchs, P.A., Elgoyhen, A.B., and Goutman, J.D. (2018). Compartmentalization of antagonistic Ca(2+) signals in developing cochlear hair cells. Proc Natl Acad Sci U S A 115(9), E2095–e2104. doi: 10.1073/pnas.1719077115.

Moglie, M.J., Wengier, D.L., Elgoyhen, A.B., and Goutman, J.D. (2020). Synaptic contributions to cochlear outer hair cell Ca<sup>2+</sup> homeostasis. bioRxiv.

Morris, G.M., Huey, R., Lindstrom, W., Sanner, M.F., Belew, R.K., Goodsell, D.S., et al. (2009). AutoDock4 and AutoDockTools4: Automated docking with selective receptor flexibility. J Comput Chem 30(16), 2785–2791. doi: 10.1002/jcc.21256.

Mukhtasimova, N., and Sine, S.M. (2013). Nicotinic receptor transduction zone: invariant arginine couples to multiple electron-rich residues. Biophys J 104(2), 355–367. doi: 10.1016/j.bpj.2012.12.013.

Mukhtasimova, N., and Sine, S.M. (2018). Full and partial agonists evoke distinct structural changes in opening the muscle acetylcholine receptor channel. J Gen Physiol 150(5), 713–729. doi: 10.1085/jgp.201711881.

Olsen, J.A., Balle, T., Gajhede, M., Ahring, P.K., and Kastrup, J.S. (2014). Molecular recognition of the neurotransmitter acetylcholine by an acetylcholine binding protein reveals determinants of binding to nicotinic acetylcholine receptors. PLoS One 9(3), e91232. doi: 10.1371/journal.pone.0091232.

Pujol, R., Lavigne-Revillard, M., and Lenoir, M. (1998). “Development of Sensory and Neural Structures in the Mammalian Cochlea,” in Development of the Auditory System, eds. E. Rubel, A. Popper & R. Fay. (New York: Springer), 146–192.

Rayes, D., De Rosa, M.J., Sine, S.M., and Bouzat, C. (2009). Number and locations of agonist binding sites required to activate homomeric Cys-loop receptors. J Neurosci 29(18), 6022–6032. doi: 10.1523/jneurosci.0627-09.2009.

Rohmann, K.N., Wersinger, E., Braude, J.P., Pyott, S.J., and Fuchs, P.A. (2015). Activation of BK and SK channels by efferent synapses on outer hair cells in high-frequency regions of the rodent cochlea. J Neurosci 35(5), 1821–1830. doi: 10.1523/jneurosci.2790-14.2015.

Rothlin, C., Verbitsky, M., Katz, E., and Elgoyhen, A. (1999). The α9 nicotinic acetylcholine receptor shares pharmacological properties with type A γ-aminobutyric acid, glycine and type 3 serotonin receptors. Molec. Pharmacol. 55, 248–254.

Rothlin, C.V., Lioudyno, M.I., Silbering, A.F., Plazas, P.V., Casati, M.E., Katz, E., et al. (2003). Direct interaction of serotonin type 3 receptor ligands with recombinant and native alpha 9 alpha 10-containing nicotinic cholinergic receptors. Mol Pharmacol 63(5), 1067–1074.

Russell, R.B., and Barton, G.J. (1992). Multiple protein sequence alignment from tertiary structure comparison: assignment of global and residue confidence levels. Proteins 14(2), 309–323. doi: 10.1002/prot.340140216.

Schwede, T., Kopp, J., Guex, N., and Peitsch, M.C. (2003). SWISS-MODEL: An automated protein homology-modeling server. Nucleic Acids Res 31(13), 3381–3385. doi: 10.1093/nar/gkg520.

Sgard, F., Charpentier, E., Bertrand, S., Walker, N., Caput, D., Graham, D., et al. (2002). A novel human nicotinic receptor subunit, α10, that confers functionality to the α9-subunit. Molec Pharmacol 61, 150–159.

Simmons, D.D. (2002). Development of the inner ear efferent system across vertebrate species. J Neurobiol 53(2), 228–250. doi: 10.1002/neu.10130 [doi].

Thompson, A.J., Lester, H.A., and Lummis, S.C. (2010). The structural basis of function in Cys-loop receptors. Q Rev Biophys 43(4), 449–499. doi: 10.1017/s0033583510000168.

Tomaselli, G.F., McLaughlin, J.T., Jurman, M.E., Hawrot, E., and Yellen, G. (1991). Mutations affecting agonist sensitivity of the nicotinic acetylcholine receptor. Biophys J 60(3), 721–727. doi: 10.1016/s0006-3495(91)82102-6.

Verbitsky, M., Rothlin, C., Katz, E., and Elgoyhen, A.B. (2000). Mixed nicotinic-muscarinic properties of the a9 nicotinic cholinergic receptor. Neuropharmacology 39, 2515–2524.

Wehrwein, E.A., Orer, H.S., and Barman, S.M. (2016). Overview of the Anatomy, Physiology, and Pharmacology of the Autonomic Nervous System. Compr Physiol 6(3), 1239–1278. doi: 10.1002/cphy.c150037.

Zhu, B.T. (2005). Mechanistic explanation for the unique pharmacologic properties of receptor partial agonists. Biomed Pharmacother 59(3), 76–89. doi: 10.1016/j.biopha.2005.01.010.

Zouridakis, M., Giastas, P., Zarkadas, E., Chroni-Tzartou, D., Bregestovski, P., and Tzartos, S.J. (2014). Crystal structures of free and antagonist-bound states of human α9 nicotinic receptor extracellular domain. Nat Struct Mol Biol 21(11), 976–980. doi: 10.1038/nsmb.2900.

